# Bayesian Metamodeling of pancreatic islet architecture and functional dynamics

**DOI:** 10.1101/2021.11.07.467656

**Authors:** Roee Lieberman, Reshef Mintz, Barak Raveh

**Affiliations:** School of Computer Science and Enginering, The Hebrew University of Jerusalem, Israel

## Abstract

The pancreatic islet (islet of Langerhans) is a mini-organ comprising several thousand endocrine cells, functioning jointly to maintain normoglycemia. Cellular networks within an islet were shown to influence its function in health and disease, but there are major gaps in our quantitative understanding of such architecture-function relations. Comprehensive modeling of an islet architecture and function requires the integration of vast amounts of information obtained through different experimental and theoretical approaches. To address this challenge, our lab has recently developed Bayesian metamodeling, a general approach for modeling complex systems by integrating heterogeneous input models. Here, we further developed metamodeling and applied it to construct a metamodel of a pancreatic islet. The metamodel relates islet architecture and function by combining a Monte-Carlo model of architecture trained on islet imaging data; and an ordinary differential equations (ODEs) mathematical model of function trained on calcium imaging, hormone imaging, and electrophysiological data. These input models are converted to a standardized statistical representation relying on Probabilistic Graphical Models; coupled by modeling their mutual relations with the physical world; and finally, harmonized through backpropagation. We validate the metamodel using existing data and use it to derive a testable hypothesis regarding the functional effect of varying intercellular connections. Since metamodeling currently requires substantial expert intervention, we also develop an automation tool for converting models to PGMs (step I) using feedforward neural networks. This automation is a first step towards automating the entire metamodeling process, working towards collaborative science through sharing of expertise, resources, data, and models.

## Introduction

### The pancreatic islet

Pancratic islets or islets of Langerhans are small micro-organs^1^ They play a major role in glucose homeostasis and systemic metabolism^2^. Due to the role of islet dysfunction and destruction in diabetes mellitus, islets are major therapeutic targets^3,4^, including the development of methods *for pancreatic islet transplantation* to treat diabetes melliitus^5-7^. The Islets are composed of multiple endocrine cell types, including insulin-secreting *β* - cells, glucagon-secreting *α* - cells and somatostatin-secreting *δ* - cells, which work together in this micro-organ to maintain normoglycemia^8^ (Fig. 1). The total number of islets in a human pancreas has been estimated to be between 3.2 and 14.8 million, with a total islet volume of 0.5 to 2.0 *cm*^*3* 9^.

**Figure 1.**
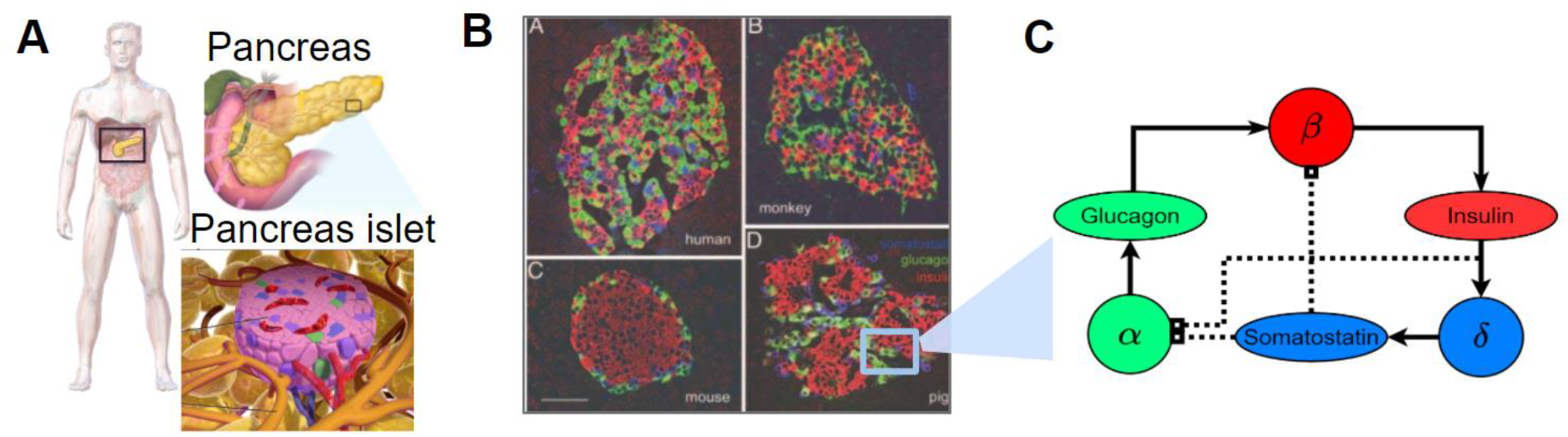
Pancreatic islet illustrations. **A** - illustration of the pancreas and pancreatic islet anatomy and tissue^43^. **B** - Islets of Langerhans show striking interspecies differences. Confocal micrographs (1-μm optical sections), showing representative immunostained pancreatic sections containing islets of Langerhans from human (*top-right*), monkey (*top-left*), mouse (*bottom-right*), and pig (*bottom-left*). Insulin-immunoreactive (red), glucagon-immunoreactive (green), and somatostatin-immunoreactive (blue) (Scale bar, 50*μm*) ^12^. **C** - *β-, α-, and δ- cell* simplify inner islet relative interaction. Straight line for excitation and dashed line for inhibition.

### Islet architecture and function

Islets have been studied extensively for decades, resulting in an established body of knowledge on pancreas cell types, but a more limited understanding of how different cell types or subtypes influence one another^9^. Although a human islet comprises only a few thousand cells, the interactions among these cells are intricate, involving electric, metabolic, and paracrine communication across many scales^1^. The cellular composition and architecture of pancreatic islets differ between and within species^8,10,11^(Fig. 1B, 2A). Such architectural differences have been previously shown to correlate with functional differences^9,12-15^ (Fig. 2B). Recent studies revealed *β - cells* subpopulations. For example, it has been suggested that islets contain *β - hub cells* or leader cells, with a higher number of intercellular connections, a unique transcriptional profile, and a functional role in coordinating glucose-stimulate insulin secretion, though their exact influence and importance are the subject of a scientific debate^9,11,16,17^. Other aspects of islet architecture-function relations are also elusive: What is the molecular basis and functional role for heterogeneity among other endocrine cells (*α-, δ-*cells)? Are subpopulations of *β* - cells (or other cells) fixed into a specific functional state, or are they transitional? ^17^.

**Figure 2.**
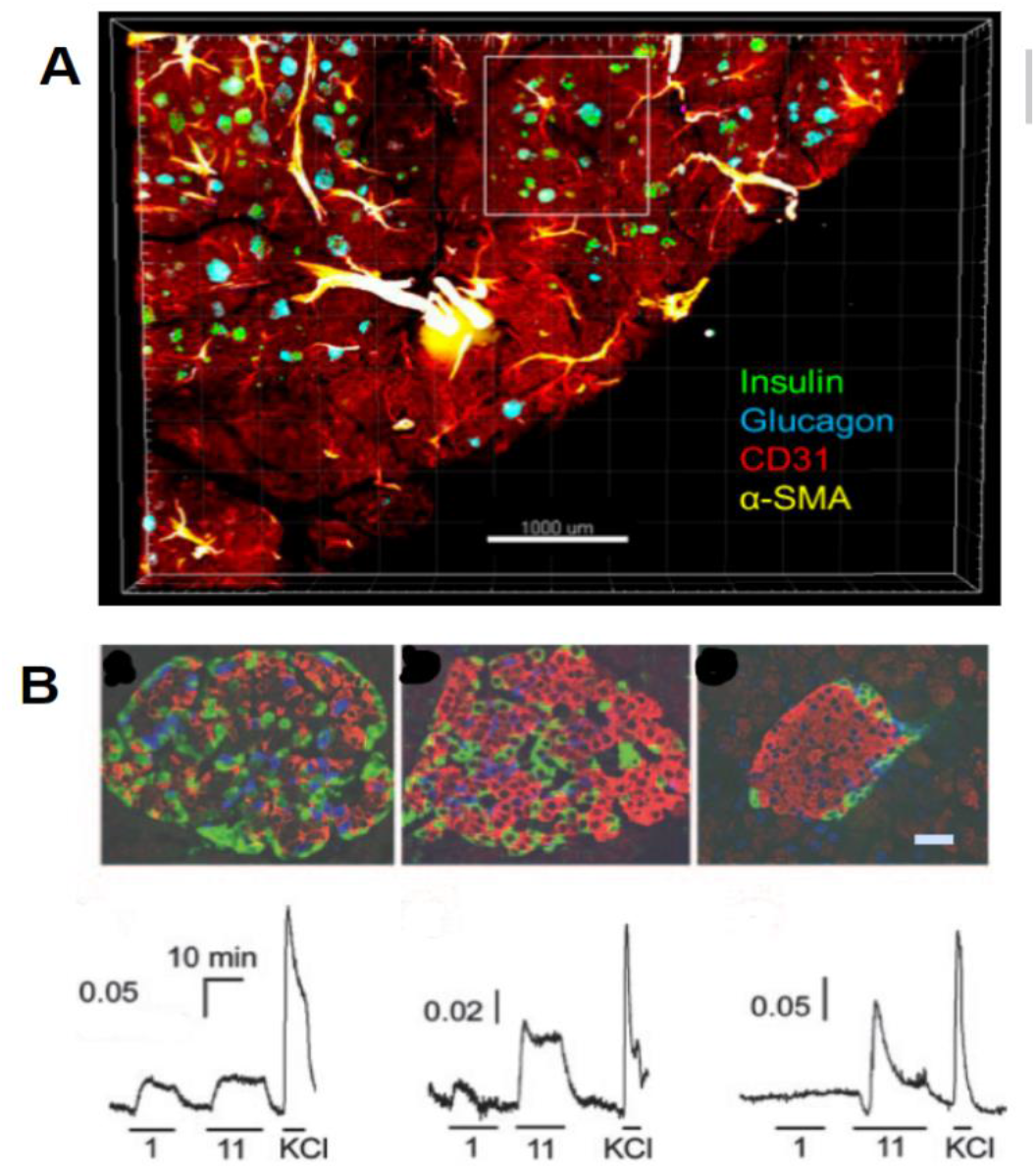
Pancreatic islet heterogeneity in architecture and function. **A** -Large-scale image capture of a human pancreas tissue slice immunostained for insulin (green), glucagon (cyan), CD31 (red), and a- SMA (yellow). Scale bar: 1,000 mm^11^. **B** -Human(left), monkey(middle), and mouse(right) islets showed functional differences that correlated with structural differences (Scale bar, 50*μm*). [*Ca*^*2+*^]_*i*_responses (Fura2-AM) elicited bylowglucose (1 mM, 1), high glucose (11 mM, 11), and high KCl (30 mM, KCl) showed that human and monkey islets, but not mouse islets responded to low glucose. Bars under the traces indicate the duration of the stimulus.^12^.

### Integrative modeling of an islet

So far, both experimental and computational tools have been used to study the above open questions^18-20^, but all of them describe only narrow aspects of an islet. A comprehensive model of a pancreatic islet is needed to further our quantitative understanding of islets and top apply this understanding to clinical use^5^. Hypothetically, we would aspire for a complete model based on the physical structure and dynamics of an entire islet, but we lack both the computational tools and the experimental data needed for computing such a model, even for a single cell ^21^ As vast amounts of data have been nonetheless collected on pancreatic islets, an integrative modeling approach could allow us to construct a useful albeit necessarily imperfect model of an islet based on as much available data as possible.

### Bayesian metamodeling

Comprehensive modeling of a whole complex system (pancreatic islet) requires the integration of vast amounts of information on various aspects of the system and its parts. To divide and conquer this task, our lab has developed Bayesian metamodeling, a general approach to modeling complex systems by integrating a collection of heterogeneous input models^21^. Each input model can in principle be based on any type of data, describe any aspect of the modeled system, and use any mathematical representation, scale, and level of granularity. These input models are (i) converted to a standardized statistical representation relying on Probabilistic Graphical Models, (ii) coupled by modeling their mutual relations with the physical world and finally, (iii) harmonized by backpropagating posterior estimates for variables in each input model in the context of all other input models ^21^. Bayesian metamodeling can in principle use any type of input model, including deterministic or stochastic, static or dynamic, and spatial or non-spatial. The only requirement is that an input model is specified quantitatively^21^. Therefore, metamodeling inherently facilitates multiscaling - coupling different input models despite significant differences in their scales. Metamodeling is also collaborative, allows autonomous contributions by different research groups with expertise spanning diverse scientific disciplines, thus maximizing flexibility, scalability, and efficiency among collaborating experimentalists and modelers^21^. However, most metamodeling procedures are currently performed *ad hoc*, using an expert to perform many steps, including conversion, and coupling, requiring a substantial amount of time and expertise. As discussed in Raveh et al. 2021^21^, the metamodeling process should be entirely automated to maximize its applicability and scalability, also providing non-experts in computational modeling with an opportunity to contribute and improve their models and the metamodel with it.

### Metamodeling of the pancreatic islet based on a new automation conversion tool

In this study, we have developed a new automated method for the conversion of input models using Bayesian networks (BNs)^22^ and feedforward neuronal networks (FNNs)^23^. We used this new tool to construct a proof-of-concept metamodel that relates the architecture and function of a pancreatic islet. The metamodel integrates two input models from the literature, each informed by different types of external data (imaging^24^, radioimmunoassay measurements^25^, etc.), based on a different modeling approach, and described using a different mathematical representation - thus demonstrating the utility of this approach for modeling a multi-cellular system. We validate the metamodel by testing its ability to reproduce known measurements and use it to generate a new testable hypothesis about structure-function relations in an islet. Our main goal is to provide a first prototype for automating the metamodeling of complex biological systems of supracellular scale.

## Results

### Definitions

We are using a few common terms that may have different definitions in different fields. Thus, we begin by defining these terms, following previously described definitions ^21^. We are working in the Bayesian framework that estimates a model based on data and prior information. Thus, a model is a joint posterior probability density function (PDF) over a set of model variables. We distinguish among three kinds of model variables: free parameters, independence variable, and dependent variables. **Free parameters** (i.e., degrees of freedom) are quantities that are fit to input information. I **Independent variables** (i.e., regressors, features, or inputs) are quantities whose values are supplied when evaluating a model. Third, **dependent variables** (i.e., response variables, outcomes, labels, predictions, or outputs) are quantities whose values are computed when evaluating a model. As an aside, **fixed parameters** (i.e., constants) are quantities whose values are defined and fixed

### Input models

The Information for the islet metamodeling is provided by two input models (Table 1) The models have been selected to describe both architectural and functional features of an islet, but neither input model describes both types of features. Both input models have been taken from published research papers based on distinct prior knowledge and experimental data on pancreatic islets. Denote the first input model as the *architecture* or *differential adhesion* model^19^, and the second input model as the *function* or *BAD* model^26^.

**Table 1:** The input models for the islet metamodeling

| Models |  |  |  |  |  |
| --- | --- | --- | --- | --- | --- |
| Model | Representation | Description | Scale | Experimental data and prior information | Ref. |
| Architecture – differential adhesion model | XYZ coordinates of balls representing islets cells | Markov-Chain Monte-Carlo sampling using the Metropolis-Hastings algorithm<sup>27,28</sup> choice to replace ball's locations and get equilibrium structure | Multicellular | Three-dimensional imaging of intact isolated islets, used to conclude the difference in the relative attraction coefficients of the Differential adhesion model. | <sup>19,29</sup> |
| Function – BAD (*beta/alpha/delta*) model | Ordinary differential equations | ODEs models for $\beta$, $\alpha$ and $\delta$ cells and their regulations on each other | Intracellular | Different molecular and electro-physiological measurements<sup>25,30</sup> to conclude the coefficients of the ODEs models | <sup>26</sup> |

### Architecture - *differential adhesion* model (Fig. 3)

The differential adhesion model computationally infers the spatial organization of different cell types in an islet from its three-dimensional islet structures based on a dataset of islet images^19^. In this work, the researchers compared to mouse, pig, and human islets, and found a conserved organization rule behind different islet structures. Based on the differential adhesion hypothesis, which postulates that differences in adhesiveness between cell types are partially responsible for the development and maintenance of organ structures^31,32^, the model is given cellular coordinates in an islet structure as input and outputs type assignments for each cell from among the alpha, beta, or delta cell types. The islet model minimized its self-energy by exchanging positions of cells, where Monte-Carlo simulation is used for the equilibration of islet structures for given cellular attractions^19,29^(Fig. 3). The conserved rule of the difference in the islet architecture between species is modeled by the relative attraction coefficients 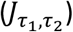 (Fig. 3, Table 2). Here, we used the coefficients that learn from three dimensional imaging of human islets^19,24^.

**Table 2:** Subset of Architecture (differential adhesion) model variables mapped in the corresponding surrogate model

| Architecture - *differential adhesion* model |  |  |  |  |
| --- | --- | --- | --- | --- |
| Free parameters |  |  |  |  |
| Name, symbol<br>*Distributed log-normal | Mean | Std-dev | Description | Unit |
| Std-dev for the dependent parameters, $\sigma$ | 0 | 0.3 | Our surrogate model uncertainty. | Connections per islet |
| Independent variables $\mathcal{X}$, $dim = 4$ | | | | |
| Name, symbol | Mean | Std-dev | Description | Unit |
| Number of $\tau$ cells, where $\tau \in \{\beta, \alpha, \delta\}$, $n_\tau$ | - | - | The number amount of the main endocrine cell of the pancreatic islet | Cells per islet |
| Number of total connections, $N_{all}$ | - | - | | Connection per islet |
| Dependent variables $\mathcal{Y}$, $dim = 6$ | | | | |
| Name, symbol<br>*Distributed normal | Mean<br>*$f$ is the FNN (Fig. 8B) | Std-dev | Description | Unit |
| Number of connections from type $\tau_1 - \tau_2$, $\tau_1, \tau_2 \in \{\beta, \alpha, \delta\}$, $N_{\tau_1\tau_2}$ | $f(\mathcal{X})$ | $\sigma$ | The number of connections from a certain type of the main endocrine cell of the pancreatic islet | Connection per islet |
| Constant parameters |  |  |  |  |
| Name, symbol | Mean | Std-dev | Description | Unit |
| Differential adhesion \ relative attractions - coefficient, $J_{\tau_1\tau_2}$, $\tau_1, \tau_2 \in \{\beta, \alpha, \delta\}$ where $J_{\alpha\alpha} = 1$ | - | - | The inferred likelihood strengths of cellular attractions, which explain the observed islet structures <sup>19</sup>. | A number |
| Temperature or fluctuation energy, T | 0.2 | - | Monte Carlo | A number |
| Neighbors' threshold factor, $ntf$ | 1.8 | - | Neighbors are cells with Euclidean distance less than the $ntf$ | A number |
| Islet's cells XYZ coordinates, $I_{XYZ}$ | - | - | The cells position in space, | XYZ cord. |

**Figure 3.**
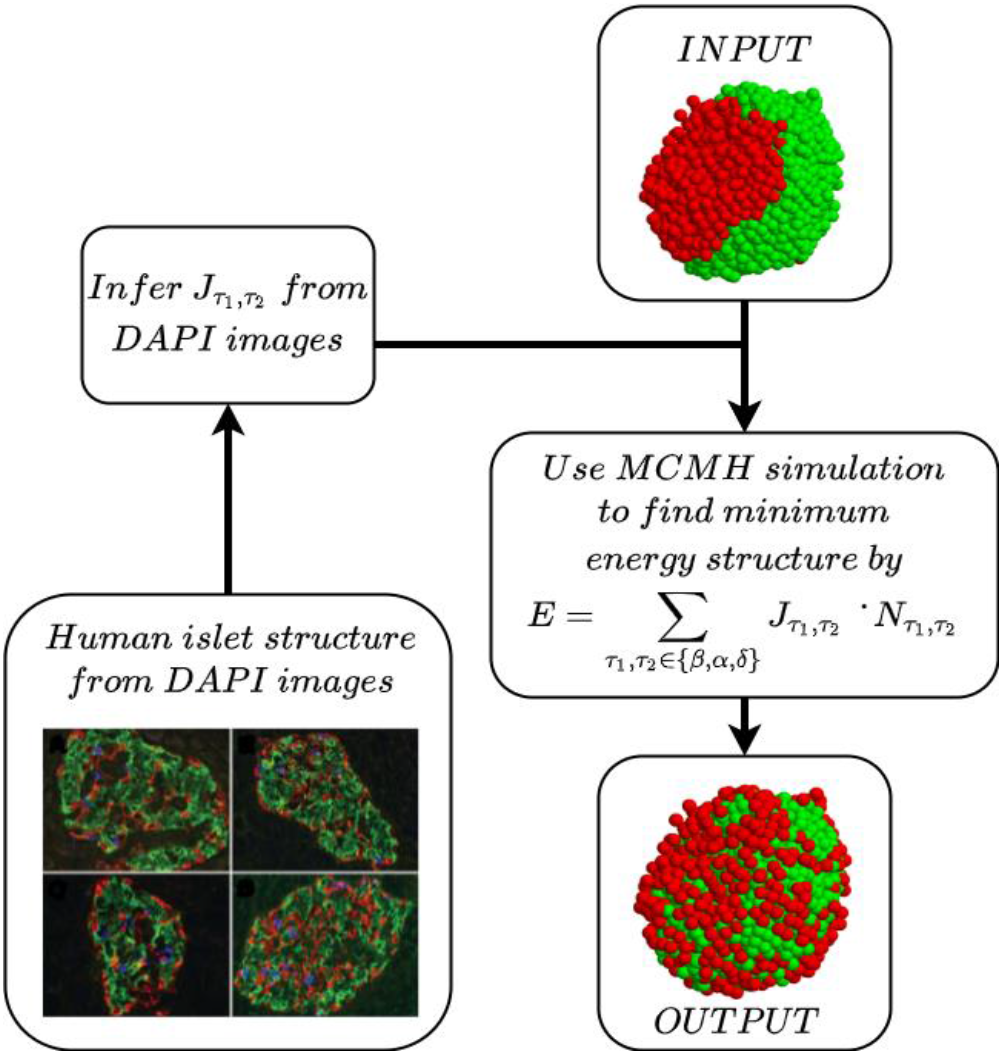
Architecture/differential adhesion model. The architecture model is given an islet architecture (<u>INPUT</u>). Using a Monte Carlo Metropolis Hasting simulation, (MCMH), it finds a minimum energy structure (<u>OUTPUT</u>). The energy function for the MCMH is based on the differential adhesion hypothesis (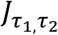 *relative adhesion strength*, 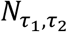*- neighbors count in islet*)(Table 2). The, 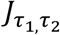 is inferred from observed islet structure from DAPI images ^19,36^.

### Function - the *BAD* model (Fig. 4)

The *BAD* model is an islet model that integrates various mathematical (ODE) models describing crosstalk among α *-*, β *-*, and δ *-* cells populations. It was developed to explore the effects of insulin and somatostatin on glucagon secretion with electrophysiological behviours^26^. The *BAD* builds on predeveloped, published ODEs models^26,33^, with new ODEs to describe the δ *- cells* and the cells interactions, based on various previous measurements (Table 1). The BAD model includes three representative cells, each describing the population mean behavior of a different cell type (*α, β, δ*) in and islet, and their averaged effects on one other over time upon glucose stimulation ^26^ (Fig. 4). The model was trained to reproduce various types of different experimental observations (patch-clamp^30^, radioimmunoassay^25^, etc.) (Table 1).

**Figure 4.**
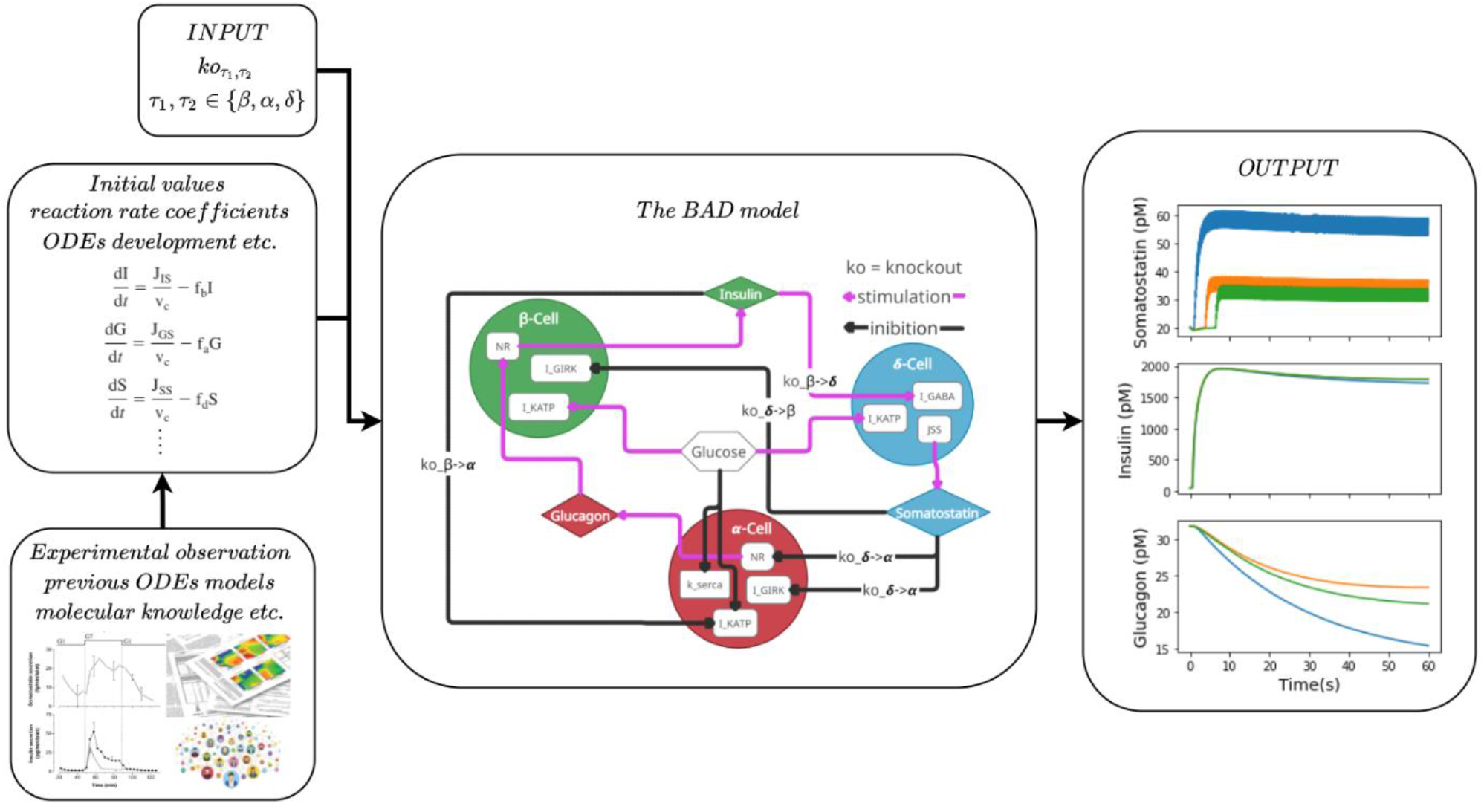
Function\*BAD* model. <u>The</u> *<u>BAD</u>*<u>model</u> is specified by a system of ODEs describing each representative cell and the crosstalk between these cells^26^. The model <u>equations and parameters (e.g. rate coefficients) were constructed</u> and fitted based on <u>previous knowledge and experimental data (e.g.</u> patch-clamp^30^, radioimmunoassay^25^). We adapted the original model to facilitate its coupling to other models through metamodeling. The <u>input</u> to the adapted model includes a connection knockout level parameter, specifying how much two different representative cells affect each other. The <u>OUTPUT</u> is a numeric estimation of key variables values over time, as illustrated for somatostatin, insulin, and glucagon over one minute following 7mM glucose stimulation.

### Conversion of input models into surrogate probabilistic models

In the first step of metamodeling, each input model is converted into a corresponding probabilistic model^21^. The purpose of this step is to transform all models onto a common representation for subsequent coupling. The probabilistic model is a surrogate for the original input model: it provides a probabilistic description of the input model, potentially simplifying it. Formally, the surrogate model specifies a PDF over some input model variables and any additional variables deemed necessary. This PDF encodes model uncertainty and statistical dependencies among its variables.

### Automating the conversion process

To partially automate the conversion process, we developed a new method that estimates the parameters of a Bayesian network (BN)^22^ using a feedforward neuronal network (FNN)^23^. For each input model, we choose a subset of variables (free parameters, independent, and dependent variables) (Table. 2, 3). To map statistical relations among the variables of each model, we reevaluate the model for different combinations of free parameters and/or independent variables values, mapped in a grid search fashion (input). For each assignment of variable values, we compute the values of dependent variables (output). We logged the values of the dependent variable as a function of the free parameters and independent variables, resulting in a matrix of observed data for each input model (Fig. 8A, bottom-left). Next, we used the observed data for learning the BN joint distribution function, with the assumption of full connectivity (Fig. 8B, 9A, 9B). To facilitate the automated conversion, we used normal distribution for the dependent variables, where the means is a function of the independent variable estimated by a simple FNN (Fig. 8B, 9) and the standard deviation is estimated by variational inference techniques from the Pyro probabilistic programming package^34^. This partial automation streamlines the conversion of arbitrary models to surrogate models compared with the formulation of the original Bayesian metamodeling method since the user only needs to specify the range of input parameters and variables assignments being evaluated (Fig 8).

### Coupling surrogate models

In the second step of metamodeling, surrogate models corresponding to multiple input models are coupled through subsets of statistically related variables. This coupling requires the introduction of some shared reference variables, which we term coupling variables Importantly, there is generally not one correct choice for how to perform the coupling step. Instead, the coupling is an external modeling choice and a model in and of itself, just like the input and surrogate models^21^(Fig 8C). In our case, to couple those two models, we took advantage of the knowledge that the *BAD* model is using a representative cell for each cell type, and that the islet regulation levels can be determined by the level of knockout 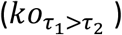 between cell type connections 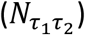. Therefore, we calculated the connections ratios 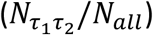 in the islet and deduced the level of knockout in a certain islet (Fig. 9C; Methods).

### Harmonize input models by backpropagation of updated variable PDFs

In the last step of metamodeling, information in the coupled surrogate models is propagated back to the original input models. This update is achieved by first updating surrogate models. We sampled from the metamodel by using uniform distribution for the model inputs (independent) variables and fit the coupling function coefficient by analyzing the results of the metamodel (Fig. 9B) (see Methods).

**Table 3:** Subset of Function (BAD) model variables mapped in the corresponding surrogate model.

| Function – *paracrine*/BAD model |  |  |  |  |
| --- | --- | --- | --- | --- |
| Free parameters |  |  |  |  |
| Name, symbol<br>*Distributed log-normal | Mean | Std-dev | Description | Unit |
| Std-dev for the dependent parameters, $\sigma$ | 0 | 0.3 | Our surrogate model uncertainty. | Connection per islet |
| Independent variables $\mathcal{X}$, $dim = 4$ | | | | |
| Name, symbol | Mean | Std-dev | Description | Unit |
| Knockout level from type $\tau \in \{(\beta \rightarrow \delta), (\beta \rightarrow \alpha), (\delta \rightarrow \beta), (\delta \rightarrow \alpha)\}$, $ko_\tau$ | - | - | The knockout level, weight for the regulation equations in the BAD model where 1 if full knockout, no regulation, and 0 is full regulation. | A number $\in [0-1]$ |
| Dependent variables $\mathcal{Y}$, $dim = 4$, the islets function features | | | | |
| Name, symbol<br>*Distributed normal | Mean<br>*$f$ is the FNN (Fig. 8B) | Std-dev | Description | Unit |
| Insulin – somatostatin threshold, $IS_{TH}$ | $f(\mathcal{X})$ | $\sigma$ | The insulin level at somatostatin bursting, | $pM$ |
| Max somatostatin, $S_{max}$ | $f(\mathcal{X})$ | $\sigma$ | The maximum somatostatin secretion level in simulation | $pM$ |
| Somatostatin secretions start time, $t_{som}^{start}$ | $f(\mathcal{X})$ | $\sigma$ | The time took to delta cell to start secreting somatostatin from the start of the simulation | $s^{-3}$ |
| Min glucagon, $G_{min}$ | $f(\mathcal{X})$ | $\sigma$ | The minimum glucagon secretion level in simulation | $pM$ |
| Constant parameters |  |  |  |  |
| Name, symbol | Mean | Std-dev | Description | Unit |
| Initial values, $I_0, G_0, \dots$ | - | - | All the initial values for the BAD model ODEs simulations<br><sup>26</sup> | Various units |
| ODE coefficients (reaction rate coefficient, etc.) | - | - | The BAD model ODEs constants<br><sup>26</sup> | Various units |

### Islet metamodel architecture-function connection

By combining the Architecture and Function model in a single metamodel, we can make quantitative predictions that can be tested through their comparison to the real world. First, we tested whether the metamodel reproduce a known empirical result. We used the metamodel to predict the suppression of glucagon secretion as connections between *β to δ*are inhibited using the connectivity knockout parameter in our metamodel (Fig. 5A). The metamodel predictions recapitulated hormone imaging data ^18^, where researchers show that *β and δ*are coupled through gap junction and use ODEs model of a whole islet that reproduce their experimental data to learn about the effect of the gap junction conductance on glucagon suppression (Fig. 5B). The second result was found when we investigate the *BAD* model. We found out that with a simple conversion rule (see methods) from the output of Model 1 (islet neighbors’ amount, 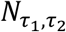) (Table 2) to Model 2 input (knockout level, *ko_τ_*) (Table 3)(Fig. 9C), we have that, different islet starts to secreting somatostatin, at a different level of insulin, and in different amounts (Fig. 6). After we accomplished the coupling, we want to find out how architecture properties influence the islets features mentioned above. We found out that the variance in those features is mostly taking place in islets with a low percentage of *β* cells (Fig. 7).

**Fig 5:**
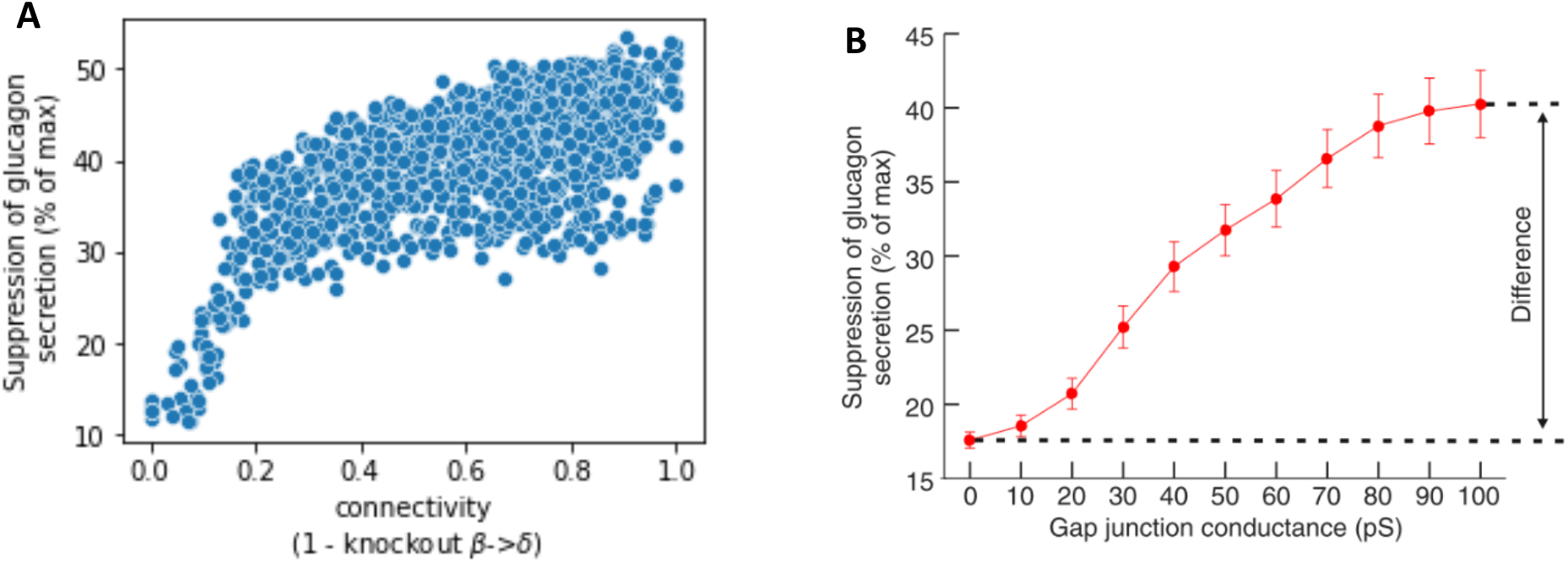
Comparison between our results to hormone imaging data measuring suppression of glucagon secretion (GlS) by high glucose (% of maximum). **A –** GlS for different values of knockout *β* → *δ* in various islet models. **B -** different values of the gap junction conductance between *β to δ* in one islet model^18^.

**Fig 6:**
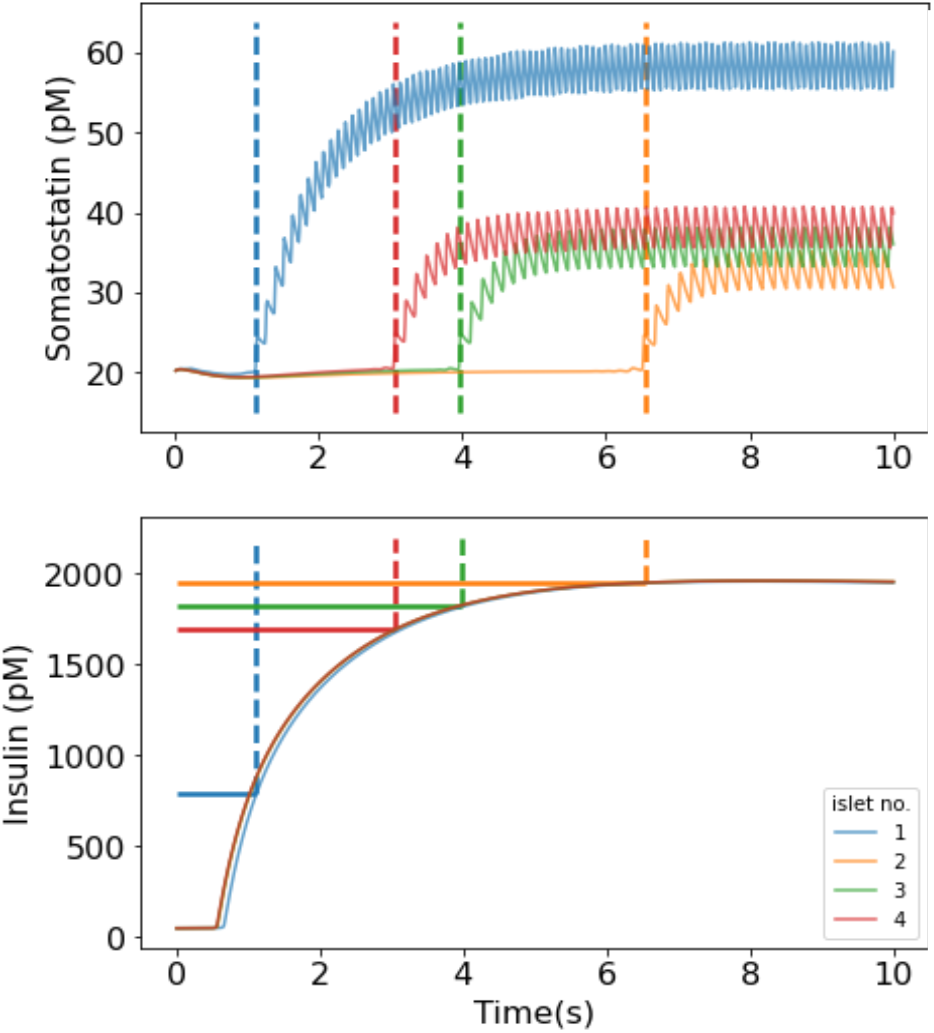
Different islet architecture start secreting somatostatin at different level of insulin and in different amount. Somatostatin and insulin secretion over time(second) in high glucose concentration, where each color is a different human islet observed architecture, These models displayed detailed morphological features based on 3D confocal reconstructions of islets ^16,19^, where islet no. This result is from the *BAD* model. We ran the model on different islet architecture in such way the architecture determines the *BAD* model knockouts levels.

**Fig 7:**
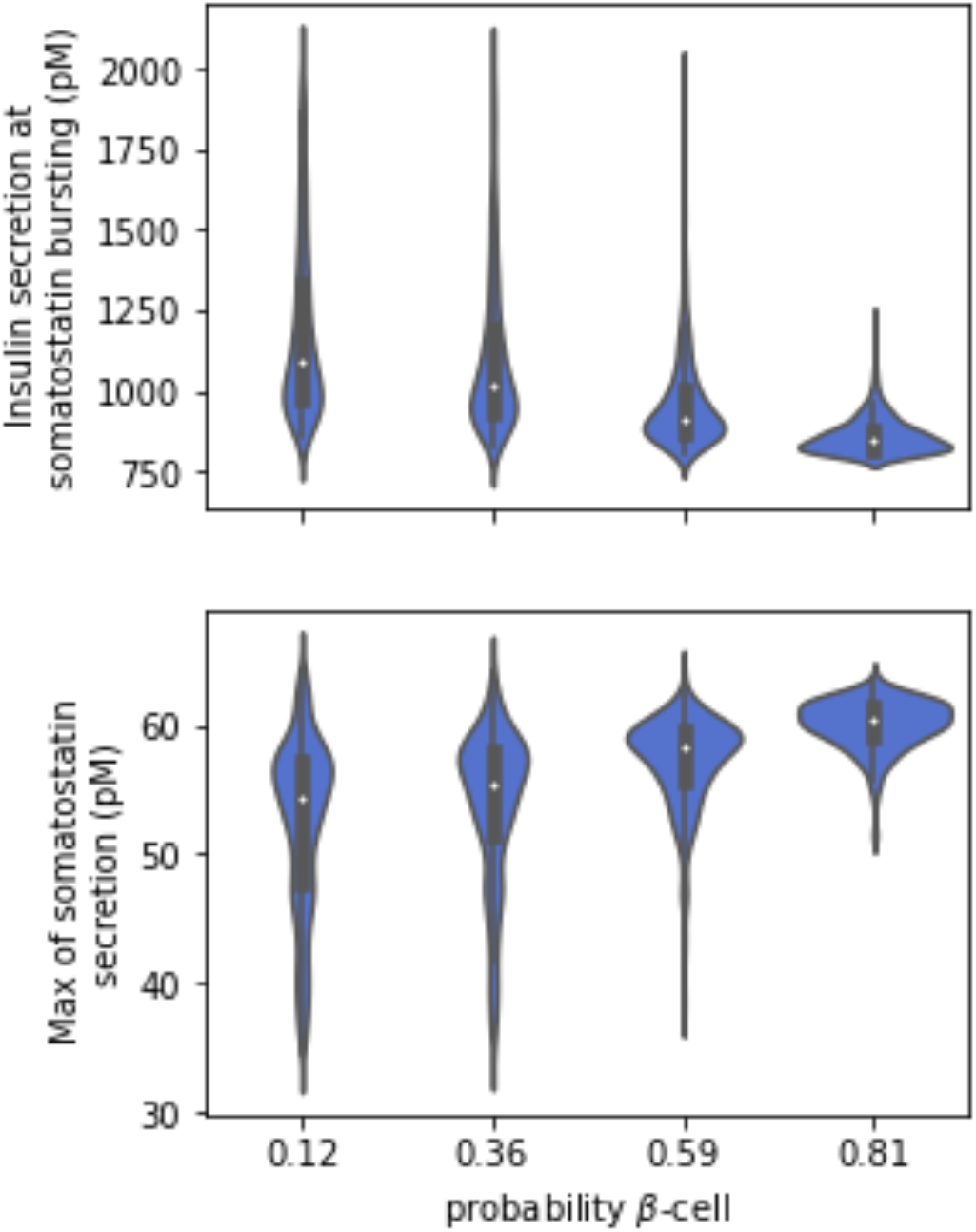
Islet metamodel sampling results. Somatostatin bursting at different level of insulin and in different amount vs different values of *β - cell* probability.

**Figure 8.**
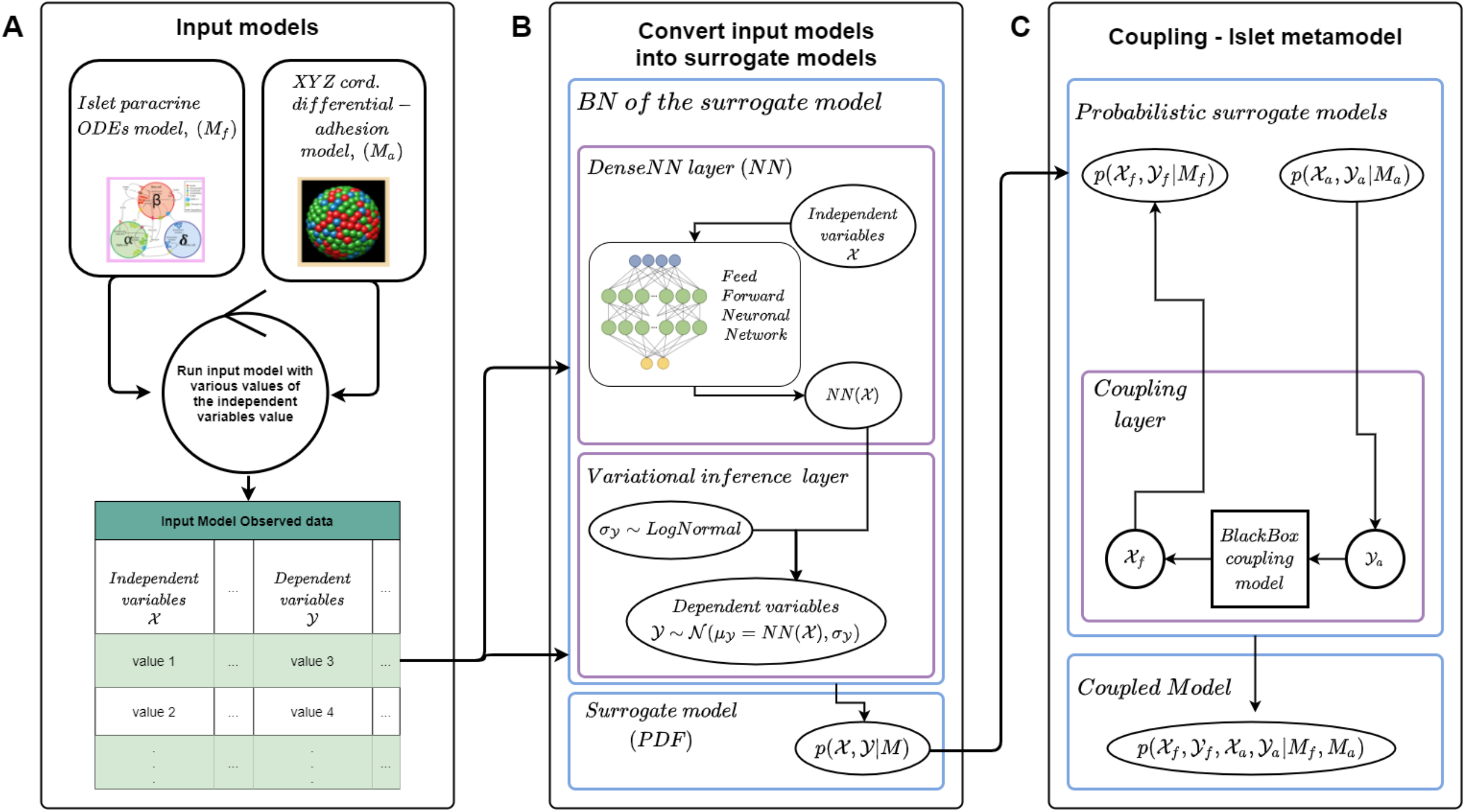
Study workflow describing the metamodeling methodology with the surrogate model converter. **A** - Run the two input models, with various values of the independent variables (and/or free parameters). **B** - Convert the models, creating a common representation, namely a PDF over the free parameters, independent variables, and dependent variable of the model using a BN and an FNN. **C** - Couple the surrogate model by statistically relating the PDFs variables using prior knowledge or some coupling model. The islet metamodel is the PDF ovel all the models’ variables.

### New testable hypothesis

Using the islet metamodel (Fig. 9) (Methods), we discover a new testable hypothesis that the variety of the insulin level at somatostatin bursting is possible only in islets with low *β* - cells ratio (Fig. 7). We saw that the insulin secretion threshold level (Fig. 6) is getting low in highly populate *β* - cells islets (Fig. 7). This behavior hasn’t been studied to our knowledge and can be tested throw experimental methods.

**Figure 9.**
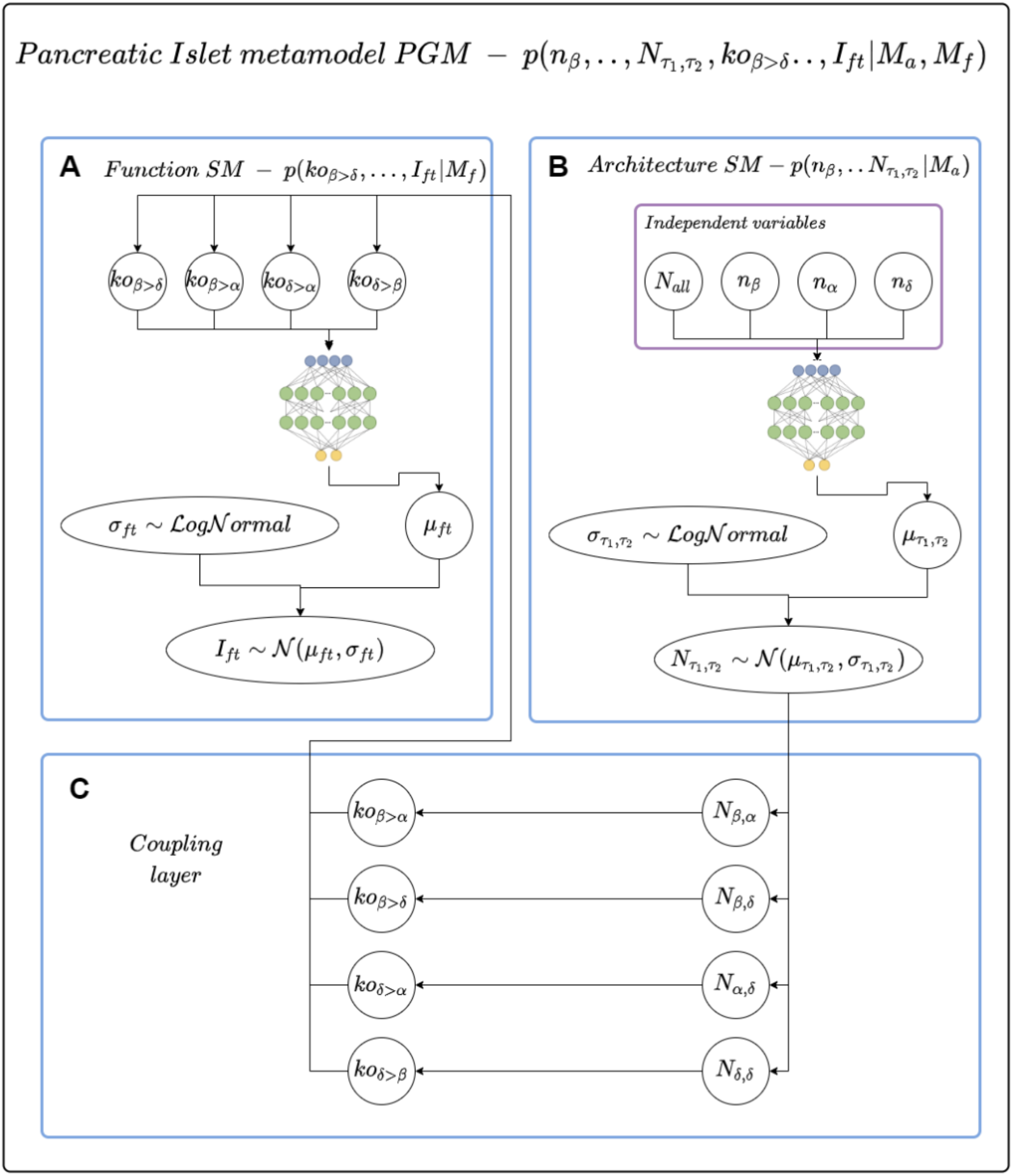
Pancreatic Islet metamodel graph (BN). **A** - The function model BN, where *ko*_*β > δ*_ stand for the knockout level of *β*-cells on *δ*-cells, where high *ko* is a low effect. The *I*_*ft*_ means islet features which is 4 features (Table 3) that we measured from the *BAD* model simulations (the dependent variables). **B** - The architecture model BN, here *n*_*τ*_ is the number of cells from type *τ ∈* {*β, α, δ*}, *N*_*all*_ is the total number of connections in the given islet structure. 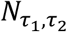 is the neighbors’ amount from type *τ*_*1*_ *- τ*_*2*_ where *τ*_*1*_, *τ*_*2*_ *∈* {*β, α, δ*}. **C** - The coupling layer we calculate the amount of *ko* based on different type of 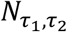 (see methods).

## Discussion

Here, we used the Bayesian metamodeling methodology^21^, a divide-and-conquer approach to modeling complex systems, in our case the pancreatic islet. We have applied the methodology for the first time on a multicellular scale system, the pancreatic islet. We combined two input models (Table 1) describing the architecture and functional features of the pancreatic islet and a coupler, to model how the architectural features affect the functional features (Fig. 9C). We also developed a new automated method based on FNNs to convert input models to surrogate models (PDF) (Fig. 8B). Finally, the resulting metamodel gave rise to a testable hypothesis about an architecture-function connection within pancreatic islets (Fig. 7).

### Combining multiple models

Combining multiple models using the same representation (i.e., the same type of model, same modeled system) is performed relatively often to increase the accuracy and precision or the coverage of the combined model. Likewise, multiple models of different types can be combined to get a model that describes a larger system and describes it more comprehensively^21^. In this study, by building a minimal comprehensive metamodel connecting the architecture of the pancreatic islet to its function, we were able to draw a new testable hypothesis. Modeling multicellular and architecturally detailed models of islet electrical activity has already been done^18^, and we show a similar result (Fig. 5), with the assumption that the regulation degree by high glucose correlates with the number of cells and the number of islet contacts.

### Drawing testable hypotheses

The pancreatic *δ*-cells, that secrets somatostatin are modulated by glucose and paracrine factors (released from neighboring *α* - cells and *β* - cells) circulating hormones and neurotransmitters released by the intra- islet nerve endings circulating hormones and neurotransmitters released by intra-islet nerve endings^15^. *δ*-cells express insulin receptors (encoded by INSR), but the action of insulin on somatostatin secretion is unclear^15^. Anterograde infusion (that is, in the direction of normal blood flow) of an anti-insulin antibody (to immunoneutralize endogenous insulin) leads to a dramatic 20-fold increase in stimulation of somatostatin secretion^18,35^. Here, our model suggests that the somatostatin bursting (delay of secretion) and the secretions’ amount, might be correlated to different insulin secretion thresholds (Fig. 6, 7). Meaning that the effect of insulin on somatostatin secretion is dependent on the islet connectivity, a rising opinion in the scientific community ^9,12,15,35^. Our model estimation is that for an islet to have a change in the somatostatin regulations the islet must have a low ratio of *β - cells* (Fig. 7). To better understand the effect of insulin on *δ- cells*, experimental tools can be used to test the hypothesis^12,18,36^. This finding might help to a better understanding and health of the pancreatic islets. As a recent review on the role of *δ- cells* in health and disease concluded: *the pancreatic islets are very complex structures and that via paracrine crosstalk, the islets are much more than the sum their parts. The δ-cells are emerging as master regulators within the islets and represent an interesting and novel pharmacological target through which dysregulated insulin and glucagon secretion in diabetes mellitus might be corrected* ^15^.

### Automating the conversion step of metamodeling

The first version of metamodeling has been applied successfully to modeling glucose-stimulated insulin secretion. However, the initial implementation had several limitations, as discussed previously ^21^. Metamodeling necessities the conversion of arbitrary models to a shared surrogate representation, and this step has thus far required manual calibration based on guessing. Here, we addressed this key limitation and developed a new simple method for the automation of the conversion step (Fig. 8B, 9). The auto converter methodology seems to work well in learning the PDF over the models’ variables. But the automation is not yet complete there are still edge cases that the current auto-converter could not handle (not normal distribution, constraints distributions, etc.), in addition, there is still a need for an expert to choose the loss functions for the auto-converter and the hyperparameters (batch size, number of epochs, learning rate, etc.) and yet the need for an expert and guessing is down significantly. Another drawback of the auto-converter is that it gives us a model that is less explanatory assuming all the dependent variables are dependent on the independent variables.

### Future application

#### Biology modeling

Because we used the metamodeling approach the current model can be easily improved given more input models or information. For example, we could add a cluster model based on single-cell RNA sequencing (sc-RNA-seq) data ^37,38^ to model the number of cells in islets and their types (the independence variable of the architecture model). That is how this minimal meta-model of the pancreatic islet can be expanded peace by peace, using the Bayesian metamodeling methodology, for accomplishing a fully comprehensive model. In addition, we can integrate this model into an already existing meta-model like the glucose-stimulated insulin secretion by the human pancreatic *β* - cell model^21^. This would facilitate exploitation of the metamodeling capability ffor summarizing information quantitatively and scalably.

#### Automation improvement

Most of the limitations of the current auto-converter can be solved with future work. There is already much work done on automated machine learning (AutoML), for learning the hyperparameters^39,40^, also there are interesting works about graphical neural networks (GNNS) that could also learn the graph structure^41^.

## Conclusion

Bayesian metamodeling is an interesting and powerful methodology that challenging the problem of modeling complex systems. With better automation, this methodology would be used by more people that could collaborate and together overcome the obstacles in the way for understanding.

## Methods

The software, input files, plots, and example output files for the present work are available at https://github.cs.huji.ac.il/ravehb-lab/islet-metamodel-pyro. For the probabilistic conversion and coupling, we used a deep universal probabilistic programming package named pyro^34^

### Extracting model observed data (Fig. 8A)

We implemented the Monte Carlo Metropolis Hasting (MCMH) simulation for the *architecture* model as described in Hoang et al., 2014^19^, and ran it 2048 times. Each simulation samples a human islet architecture from the 12 islets given in Hoang et al., 2014^19^, that can be found in /https://github.com/chonlei/bHub_sim^16^, then randomly chose types for the cells from *p*_*β*_, *p*_*α*_, *p*_*δ*_ *∼ Dirichlet* (1,1,1) and ran the MCMH simulation for one million steps each. For the *BAD* model ODEs, we use the same tools as the in BAD model paper^26^. we use the initial values and the equation that supplies https://github.com/artielbm/artielbm.github.io/tree/master/Models/BAD^26^, Figure6.ode, and ran the model using the XPPXUT software^42^ as described in the paper. We simulated for 5 minutes, with 32 resolution ranges range on all the ko types (Table 3) permutation. The chosen variables and parameters we looked at can be found in the supplied information (Table 2, 3).

### Using the surrogate model auto converter

The Jupiter notebook with all the train parameters for the conversion of the input model can be found in the git directory (link above)

### Coupling

The coupling between the two surrogate models was done by relating the neighbor’s ratio variable 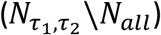 in Model 1 (Fig. 9B, C, Table 2) to the knockout parameter (*ko*_*τ*_) in Model 2 (Fig. 9A, C, Table 3) in the following formula:

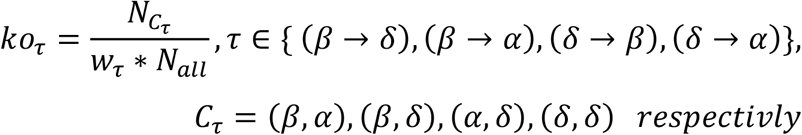

The values of *w*_*τ*_ (coefficients for the equations) can be found in the attached git directory (link above). The values of *w*_*τ*_ was fitted to stretch the neighbor’s ratio 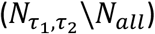 variables in between 0 and 1. The rationale for this coupling scheme is based on the finding that the islet regulation is affected by its connections^18^, and from recent findings that show that *δ*-cells are dynamics and can send arms in order to change their connectivity and relativity to other cells types^20^.

### Sampling from the prior uniform distribution of the input variables

To estimates the islet meta-model, we sample the left independent variable (Fig. 9B) from a uniform distribution and ran it throw the metamodel, and examine the results (Fig. 5, 7).

## Notes

https://github.cs.huji.ac.il/ravehb-lab/islet-metamodel-pyro

### Competing Interest Statement

The authors have declared no competing interest.

